# Neurodiversity in the brain: More variable localization of face regions in autism

**DOI:** 10.64898/2026.05.14.724682

**Authors:** Amanda M. O’Brien, Tyler K. Perrachione, John D. E. Gabrieli, Anila M. D’Mello

## Abstract

A fundamental principle in human neuroscience is that the brain is organized into distinct functional regions specialized for particular processes. These functional regions develop early and are shaped by sensory experiences. Strikingly, the precise location of these functional regions is relatively consistent across individuals, with specific regions found in stereotyped locations with respect to the macroanatomy (e.g., particular sulci and gyri). An important question is how flexible this functional neuroanatomy is, particularly in neurodevelopmental disorders like autism which are characterized by atypical brain development and behavior. Here, we investigated this question by focusing on the organization of the face-processing network — a well-characterized system of regions that are reliably localized across neurotypical individuals, including, famously, the fusiform face area. Using precision fMRI, we identified subject-specific face-sensitive regions in the brains of autistic and neurotypical adults, and assessed their topographical alignment across individuals. Face regions in autism were globally and highly variably displaced – located farther from their expected locations in the brain relative to much more homogenous localization in neurotypical adults. Autistic individuals whose face regions were more displaced had greater sociocommunicative challenges. In contrast, the spatial organization of object-sensitive regions was not affected. These findings suggest that the spatial organization of the face network is atypical in autism, with behaviorally meaningful increased variability in the precise location of face-sensitive areas, and highlights the importance of individualized approaches in neuroimaging.

**Significance Statement:** The human brain is organized into specialized functional regions positioned in remarkably consistent locations across individuals. It is unclear how altered neurodevelopment affects this organization. Using precision fMRI, we found that autism was characterized by globally and highly variably displaced face-sensitive functional brain regions. Atypical spatial organization of these regions was associated with greater sociocommunicative difficulties. In contrast, the spatial organization of object-sensitive regions did not differ in autism. These findings reveal that functional neuroanatomy in autism is altered in category-specific ways, with direct relevance to core sociocommunicative features of the condition. This work underscores the importance of individualized brain mapping in neuroimaging and may shed light on inconsistent findings in previous group-level neuroimaging studies.

---

A fundamental principle in human neuroscience is that the brain is organized into distinct functional regions specialized for particular cognitive, sensory, and motor processes (i.e., the ‘functional neuroanatomy’ of the brain). These functional regions are found throughout the cerebral cortex and are associated with particular macroanatomical locations (1, 2). In the domain of visual perception, several proximal but discrete functional regions are distributed throughout the cortex, forming networks of regions specialized for processing distinct visual categories (3, 4). Perhaps the most famous example is the face processing network, comprised of several occipito-parietal and frontal regions engaged by face processing. Importantly, individual face processing regions including the fusiform face area (FFA) - a region specialized for the early identification of faces (5) - are highly spatially constrained. Indeed, the location of the FFA is largely overlapping and consistent across people (6). There are nonetheless individual differences in the precise spatial location of functional regions, which may be behaviorally meaningful (for evidence for the link between spatial location and behavior in other domains, see (7). Neural responses extracted from individually-defined functional regions, such as the FFA, can be strikingly different from those extracted from group-averaged data and lead to different conclusions about the selectivity or function of this region (8). This suggests that individual differences in *topography* or spatial organization of functional regions is an important consideration when investigating brain function (9), and that group-level approaches that spatially average across individuals could obscure meaningful results.

Consideration of individual differences in functional neuroanatomy are particularly relevant when investigating the brain in neurodevelopmental conditions such as autism spectrum disorder (ASD). ASD is a spectrum-based condition characterized by substantial behavioral *heterogeneity -* meaning that behavioral patterns (e.g., visual attention, gaze patterns, etc.) are less consistent across ASD individuals than neurotypical (NT) individuals. For instance, while NT individuals show high interindividual consistency in their interest and attention to faces, such as focusing on the eyes and mouth, looking patterns across autistic individuals are highly variable (10, 11). This behavioral heterogeneity is mirrored at the neural level. Faces are a visual category which is typically strongly and consistently represented in specific regions of the brain in neurotypical individuals (e.g., the occipital face area, FFA, medial prefrontal cortex, inferior frontal gyrus; (12). However, neuroimaging studies indicate that both whole-brain functional network organization and face-evoked responses are more variable in autistic individuals (13). This heterogeneity complicates assumptions that underlie group-averaging neuroimaging approaches. It is possible that individual differences in the organization of face regions in autism contribute to heterogeneous behavioral and neural responses to faces. Despite the growing adoption of precision neuroimaging methods (14), few if any studies have investigated individual differences in the functional neuroanatomy of autism.

In the present study, we quantified the (neuro)diversity of functional neuroanatomy of face processing in autism (vs. neurotypical brains). In contrast to a group-averaging approach, we used a *precision-fMRI* analysis: identifying functional regions sensitive to faces *in each individual* to investigate functional neuroanatomy. We further probed whether any differences in autistic functional neuroanatomy were (1) more prominent in specific face regions associated with early (posterior) versus late (frontal) perceptual processing; (2) related to individual differences in autism-relevant behaviors; and (3) stimulus specific (e.g., present only for faces or also for other visual categories such as objects).

## Results

### Identification of individually defined face regions

We used an established fMRI repetition suppression task known to robustly distinguish face processing regions (15, 16). In all individuals, the target contrast (non-repeating > repeating faces) activated the canonical face network including the bilateral fusiform gyrus (housing the FFA), occipital face area, posterior parietal, inferior frontal gyrus, and medial prefrontal cortex (**Figure 1A**). We next created data-driven parcels, delineating areas of high probability of activation across participants (**Figure 1B**). These parcels overlapped with the canonical face processing network, capturing activation localized both in posterior regions (such as the fusiform face area, occipital face area, and posterior parietal cortex) that support early identification and perception of faces, as well as in prefrontal areas (such as the mPFC and inferior frontal gyrus) that support higher-level face processing including face memory and dynamic and emotional face processing (17). We used these parcels to constrain the identification of individually defined functional regions of interest (“face regions”) in each participant (18), comprising the top 10% most active voxels in each parcel (**Figure 1C**). The spatial organization of, and functional responses within, these face regions were probed in subsequent analyses.

**Figure 1.**
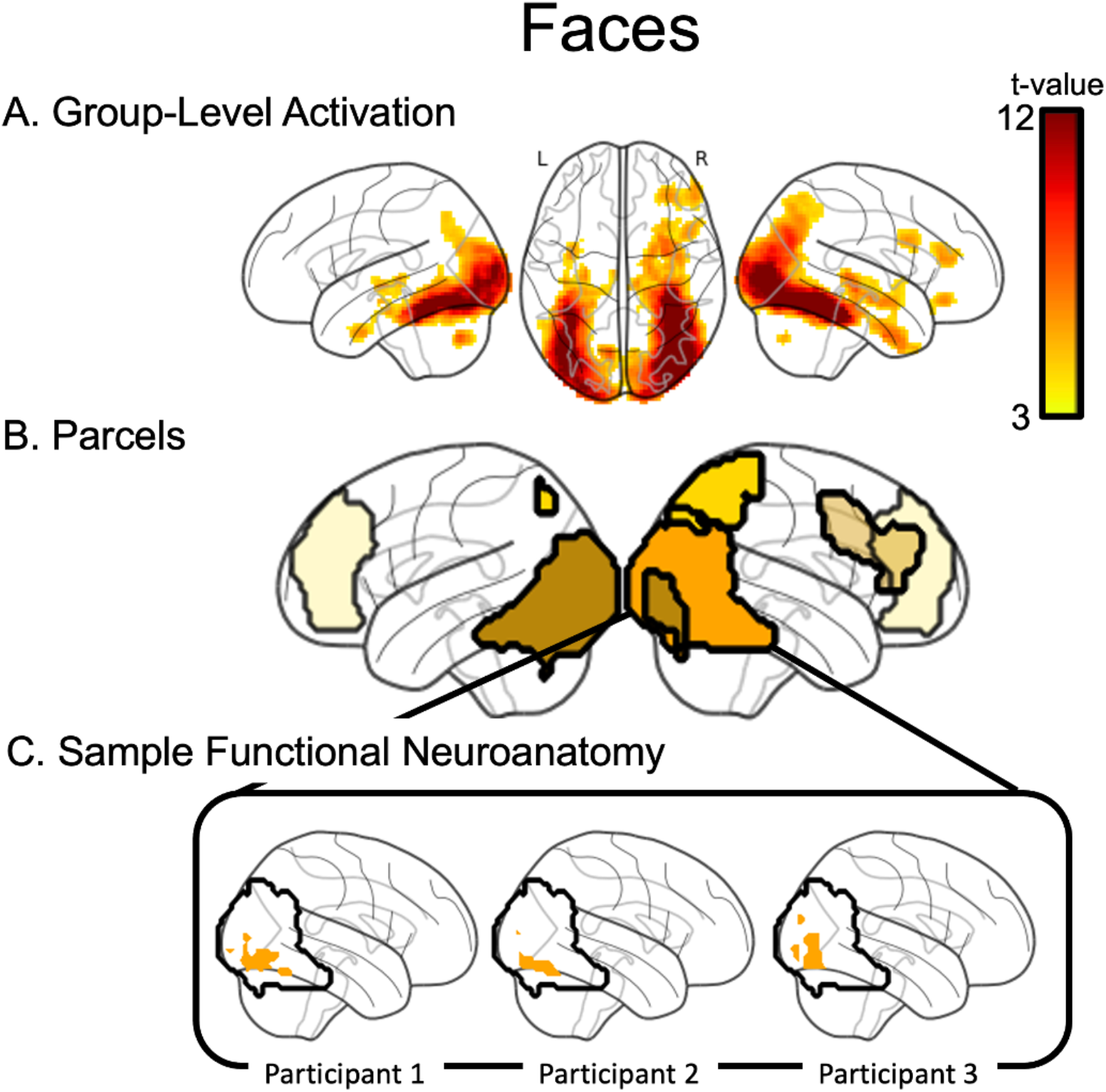
Functional tasks activate face-specific functional regions. **A**. Group-level activation across all participants for faces, thresholded at *p*<0.001, FWE-corrected. **B**. Five functional data-driven parcels were identified that capture areas of high probability of face-specific activation across participants (Methods 3.3). **C**. Individually-defined functional face regions (top 10% of voxels within the parcel) from three example participants for the parcel including the FFA (Methods 3.4).

### Displacement of individually defined face regions from canonical locations

We first examined the spatial displacement of each participant’s individually defined face regions relative to standard reference locations in the brain (see **Figure 2A** for example; See **SI-Table 1** for coordinates of reference locations). For each participant, we calculated the mean Euclidean distance between every voxel in their face region and a functionally defined centroid of activation for each parcel based on the center of mass of the face regions in the NT group. This quantifies the degree of spatial deviation for each individual from the expected location for that parcel. We examined whether the amount of displacement differed between NT and ASD (effect of Group) and between posterior and frontal parcels (effect of Location).

**Figure 2.**
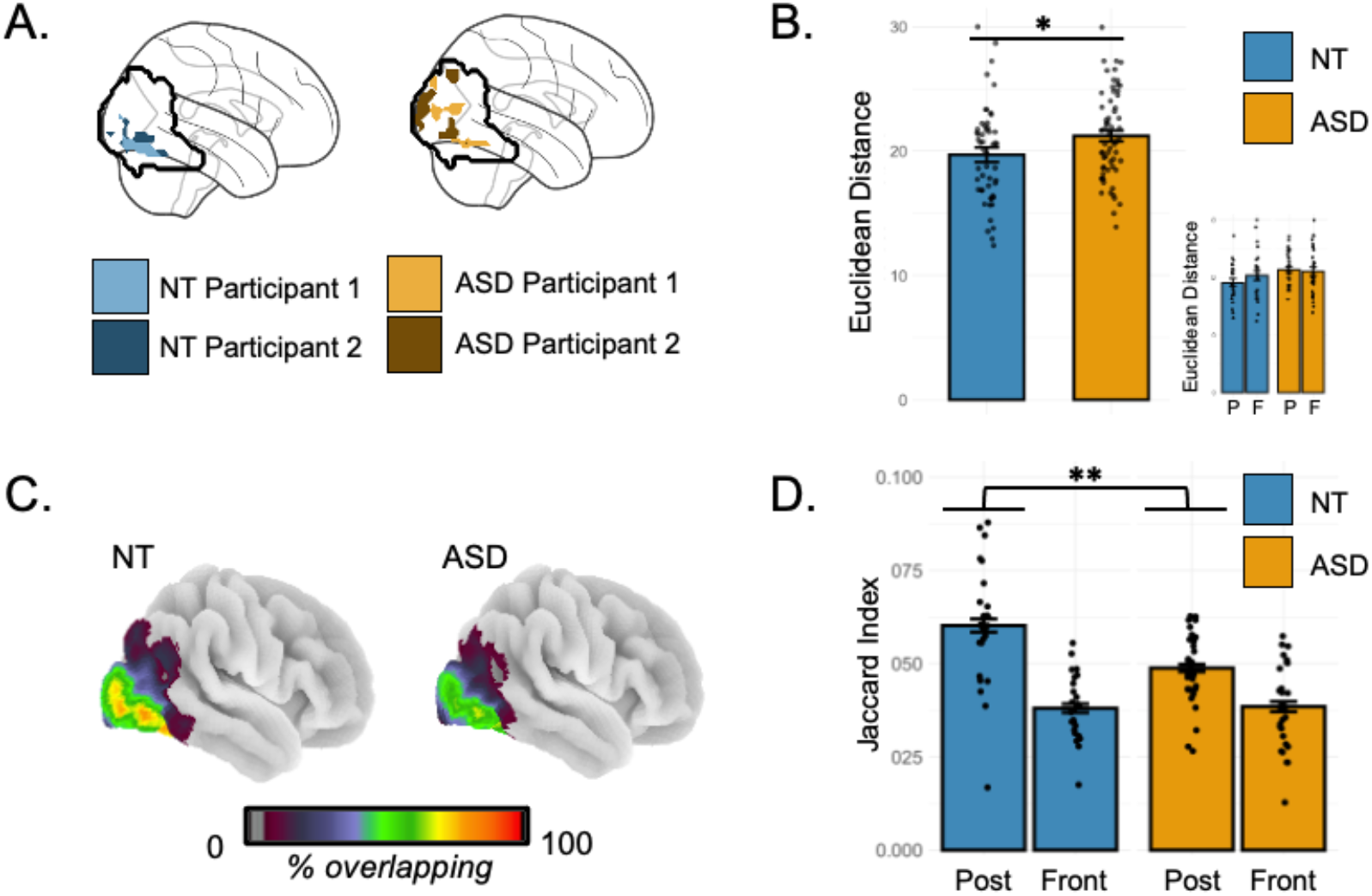
Atypical organization of face-specific functional regions in ASD. **A**. Example of spatial overlap of two participants’ functional regions by group in one parcel. **B**. Autistic individuals showed increased displacement of functional regions from the expected location (as measured by Euclidean distance between each participant’s functional region and an NT centroid). **Inset**: Displacement further divided by region (posterior, frontal). **C**. Voxel-wise overlap (i.e., probability maps) showing the percentage of people that show overlapping activation at each voxel in one example parcel. **D**. There was a significant interaction between diagnosis and brain location (posterior, frontal). The organization of functional regions in the posterior brain (associated with earlier stimulus processing) was more variable in ASD than in NT individuals. Note: Dots represent individual participants. Error bars represent standard error. Yellow=ASD, Blue=NT. P/Post = Poster face regions; F/Front = Frontal face regions.

Individually defined face regions in autistic individuals were located farther from their expected locations (**Figure 2B;** mean absolute distance in mm; main effect of Group, *F*(1,58)=4.50, *p*=.038). There was no main effect of Location and no Group × Location interaction, indicating a global shift in face regions in ASD away from their expected or canonical locations in the brain.

### Dispersion in the locations of individually defined face regions in autism

Given the global displacement of face regions in the ASD group, a follow-up question is whether ASD individuals collectively rely on a different yet consistent set of face regions (i.e., an “ASD-specific” face network), or whether this displacement reflects greater heterogeneity in the location of face regions across ASD individuals. To investigate this, we quantified the spatial dispersion of face regions within each group by measuring the voxel-wise overlap between each participant’s individually defined face regions and those of all other participants in their group.

There was greater spatial dispersion of face regions in ASD individuals than NT individuals (**Figure 2C**), an effect that was stronger in the posterior than frontal regions (Group × Location interaction, (*F*(1,223)=10.52, *p*=.001; **Figure 2D**). This suggests that autism is characterized by greater heterogeneity in the location of face processing regions important for the earliest stages of face perception.

### Impact of functional region size on dispersion

One possibility is that the finding of increased dispersion of posterior face regions in autism was affected by the fROI-forming threshold. For instance, defining regions as the top 10% of activated voxels may have been overly conservative in estimating the extent of each autistic individual’s face-sensitive areas; coupled with the possibility that functional responses may be sparser in autism than NTs, constraining the definition of a functional region to just the top 10% of activated voxels might inadvertently underestimate the degree of overlap across ASD participants. We therefore investigated whether/how group differences in dispersion changed with three increasingly liberal thresholds for face-region inclusion (top 20%, 30%, or 40% of the most activated voxels, yielding progressively larger fROIs). Autistic individuals continued to show greater dispersion than NTs even when defining face regions to include larger swaths of cortex (**Figure 3**).

**Figure 3.**
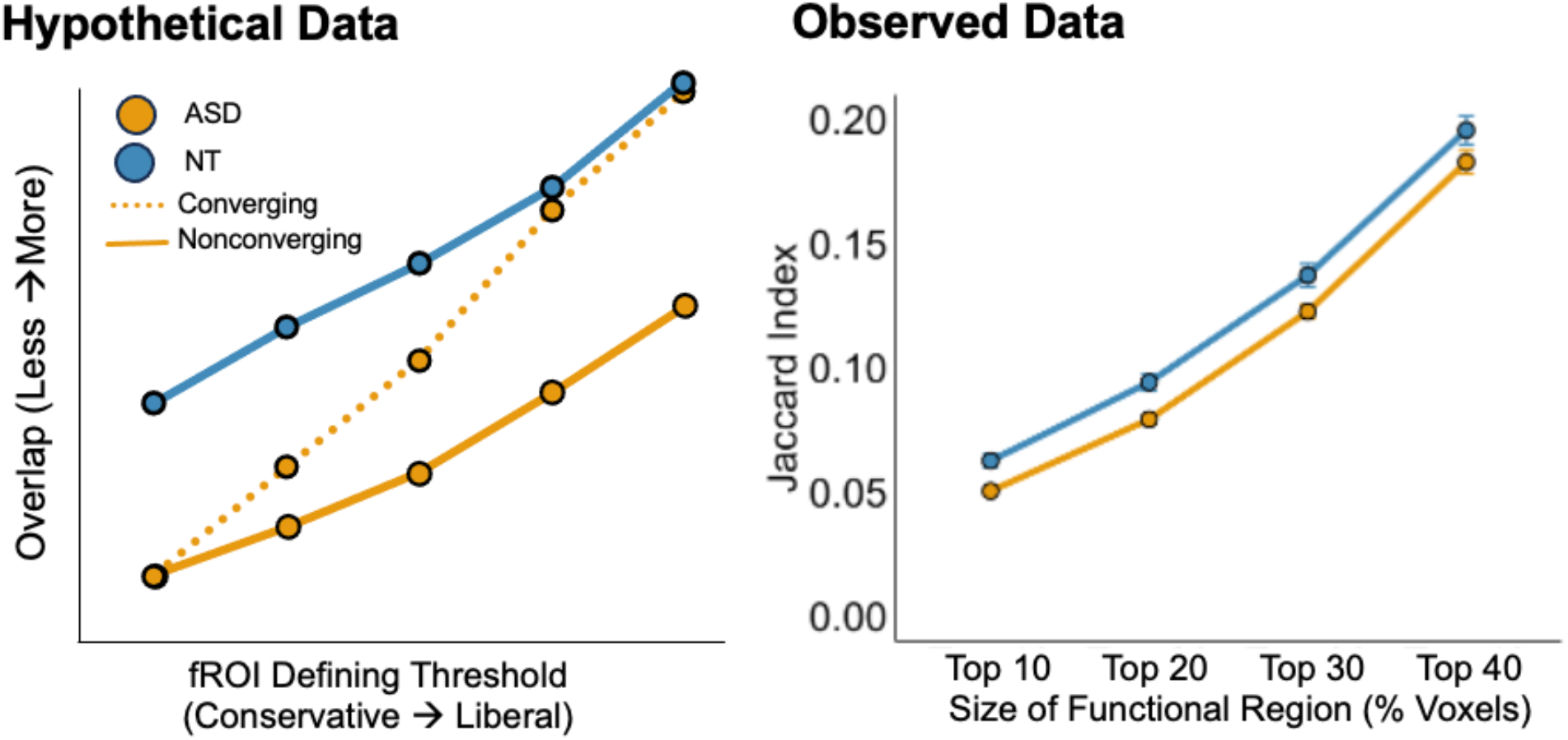
Increased dispersion of functional regions in the autism group is observed in posterior regions even when region size is increased (top 20%, 30% and 40% of voxels in a given parcel). **Left**. Hypothetical data illustrating potential outcomes of the analysis. The dotted line shows an outcome in which face regions in ASD are more spatially dispersed than in NT individuals when considering the top 10% of voxels, but converge on NT levels of dispersion when face regions are defined more liberally. The solid line shows an outcome in which the dispersion of face regions in ASD is consistently larger than the NT group, regardless of inclusion threshold used to define the region. **Right**. Observed data show that dispersion of face regions was consistently greater in ASD even across increasing region size.

### Associations between displacement of functional regions and autism symptoms

Are individual differences in functional neuroanatomy *behaviorally* meaningful? To investigate this, we correlated the degree of face-region displacement with autism symptom severity, as measured by the ADOS Total Score. We found that greater displacement of face regions from their typical location was associated with greater autism symptomatology. This effect was specific to posterior regions and insignificant in frontal regions (posterior r=0.44, *p*=.016; frontal r=0.10, *p*=.262) (**Figure 4**).

**Figure 4.**
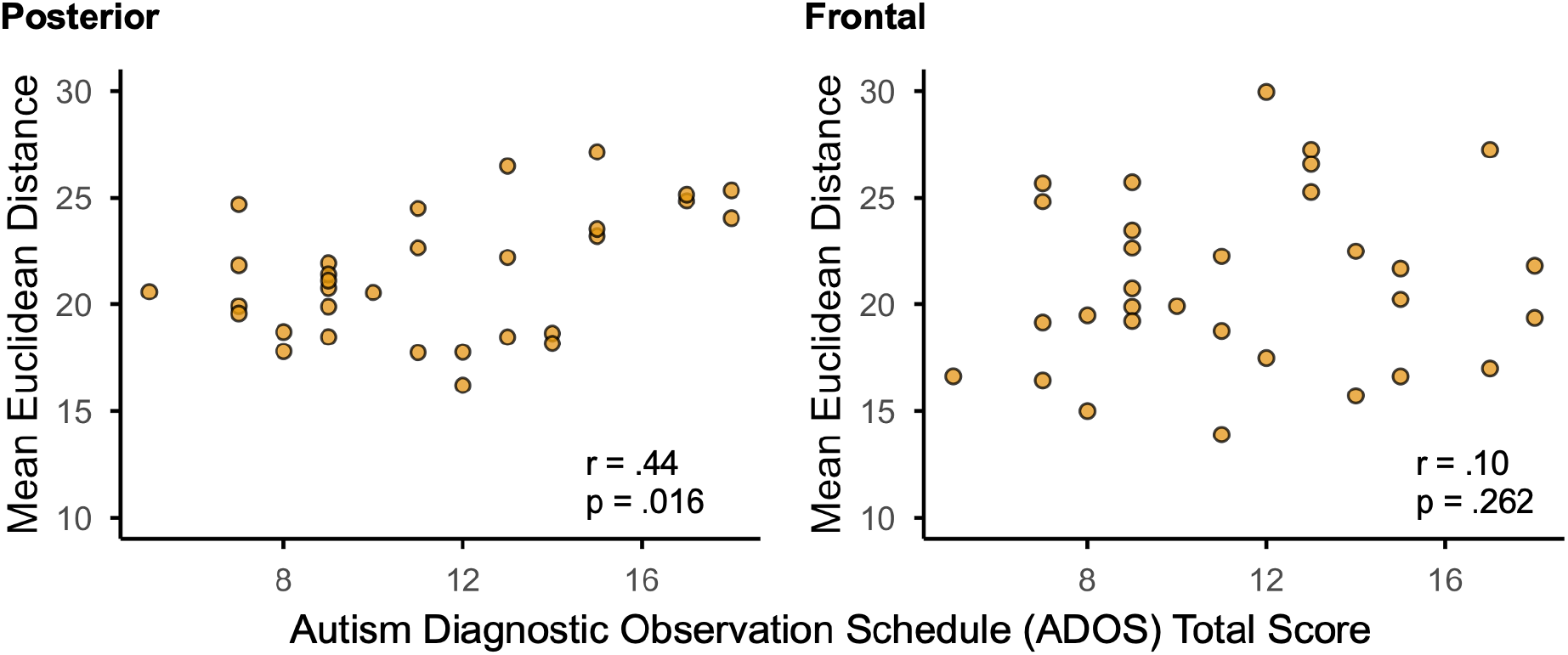
Spearman’s Rank Correlations between displacement measure (how far face regions were located from their expected location in the brain) and autism traits.

### Category-specific group differences in functional neuroanatomy

A critical question is whether the atypical functional neuroanatomy we identified is specific to face-processing, or whether it is also evident in the organization of other visual functional regions, such as those that process objects. We conducted a similar set of analyses on the organization of *object*-specific functional regions (See **SI-Figure 1** for visualization of functional object-regions). Unlike faces, we found no group differences in displacement or dispersion of object-sensitive regions, and no associations with behavior (all p’s > .05; See **SI-Table 2**). This suggests that the greatest atypicality in functional neuroanatomy in ASD is found in domains particularly relevant to or most affected in autism (e.g., face processing).

### Potential confounds contributing to atypical functional neuroanatomy

Atypical functional neuroanatomy could be driven by a number of confounds including noisier data (i.e., reduced consistency in magnitude of activation over time) or group-wise magnitude differences (i.e., ASD individuals could show overall lower activation in their own face regions, resulting in the appearance of reduced overlap). We found no effects of group related to noisiness of fMRI data (defined as run-to-run variability in voxel-wise response profiles across regions, ASD: *r*=.30, NT: *r*=.31, *t*(54)=-0.04, *p*=.97). There were also no group differences in parcel-wise activation magnitude (*t*(127)=0.67, *p=*.503). Finally, there were no group differences in the response magnitude of individually defined face regions to the non-preferred stimulus category (objects) (*t*(87.5)= 1.25, *p*=.22). This provides evidence that the functionally defined face regions were not engaged in categorically different processing between groups.

## Discussion

The human cerebral cortex consists of a mosaic of distinct functional areas (19). The spatial location of these functional areas is a key determinant of their function. While the macroanatomical location of these regions is generally consistent across individuals, there is variation in their precise location across people, which has been associated with individual differences in behavior. Here, we show that autistic individuals exhibit a brain-wide displacement of face-sensitive regions relative to their canonical locations. Greater displacement of face-sensitive regions from their expected locations in the brain was associated with greater difficulty in social communication in autism. In addition to global displacement of face-sensitive regions, we observed greater spatial dispersion of face-sensitive regions in autism in posterior occipito-parietal cortex (see **Figure 5A** for graphic summary), which plays a critical role in the earliest stages of face perception. Interestingly these effects were specific to the face-processing system; we did not find similar atypical spatial organization of object-sensitive regions.

**Figure 5.**
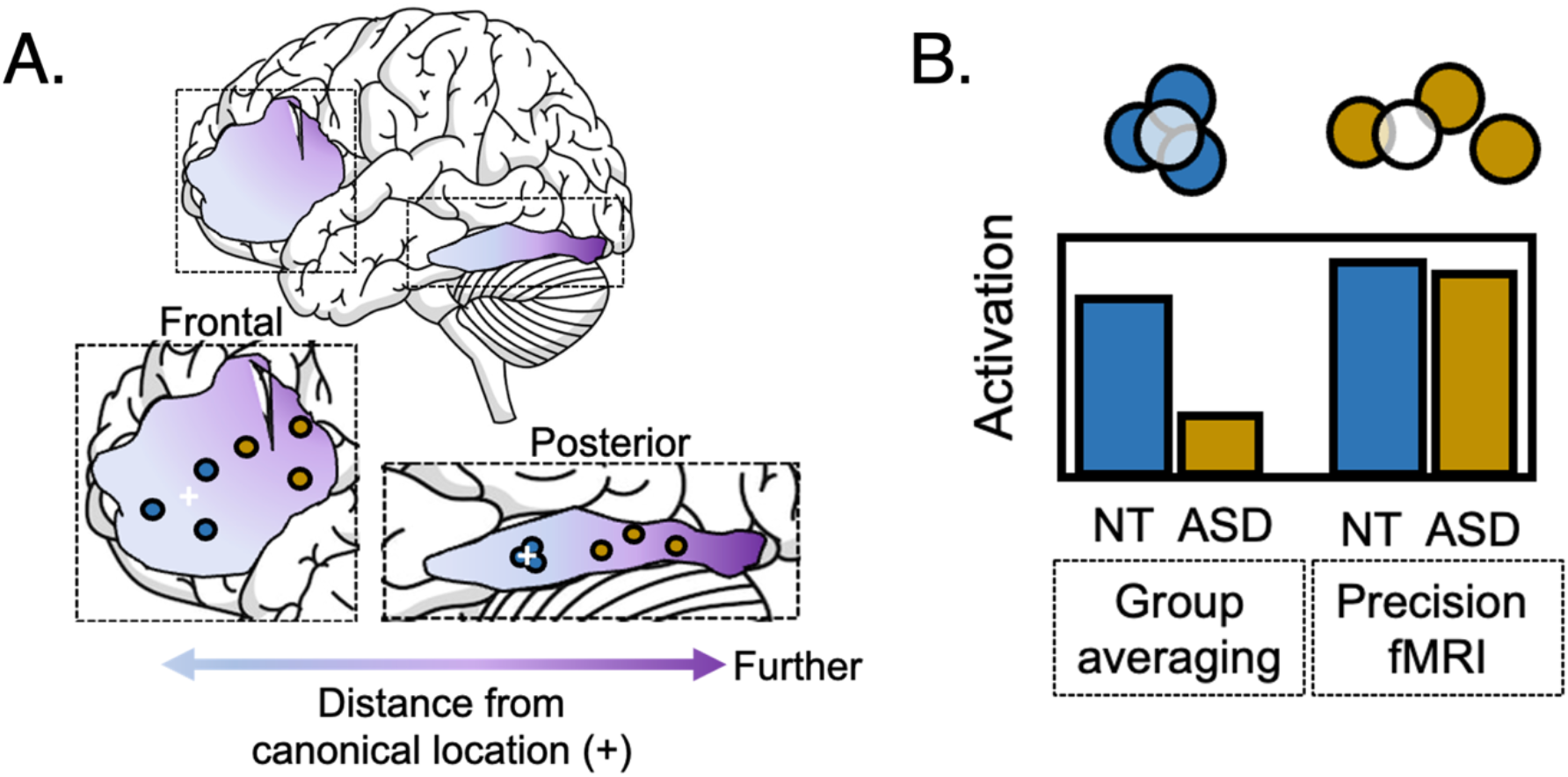
Illustrative figures and impact on neuroimaging. **A**. Schematic of results in frontal and posterior areas. Across posterior and frontal parcels, individually defined functional regions in autism (represented by yellow dots) were located farther away from their canonical location (represented by light-dark purple spectrum and white +). In posterior parcels, NTs (blue dots) showed high consistency in the location of functional regions while ASD individuals showed heterogeneity. In frontal parcels, both ASD and NT individuals showed heterogeneity in the location of individually defined functional regions. **B**. Illustration of potential impact of increased spatial variability when traditional group-averaging methods are used. The yellow and blue dots represent the hypothetical location of activation from individual autistic and NT participants respectively, and the white circle represents a small region of interest used for extracting average group-level activation. Taking the average of the NT activation from the black circle captures the responses of every single NT participant. In contrast, taking the average activation from the same location in the autism group only captures the activation from a fraction of the autistic participants. Even though all autistic participants are indeed showing activation in response to the stimuli, this ‘displaced’ activation is largely not captured when averaging across individuals within a small region of interest. This results in apparent reduction of activation in the autism group relative to precision fMRI approaches which consider each individual’s peak activation, as illustrated by the hypothetical bar chart.

### The location of functional regions matters

While it is widely appreciated that the human brain comprises diverse functional regions that tile the cortex, only recently has there been greater recognition of substantial individual variability in the precise location of these regions. Most neuroimaging studies have relied on group-averaging approaches, which spatially blur and conflate functionally distinct regions (20), potentially obscuring meaningful individual differences. Regardless of individual differences, the precise spatial location of functional regions in the brain may be significant - constraining the functions a given region supports. Functional regions are situated within cytoarchitectonic contexts that are well suited for particular computations (21) and are optimally positioned within broader networks to receive the appropriate inputs from other brain areas. This suggests that even slight alterations in the location of functional regions may affect function. Indeed, the location of an individual’s functional networks is highly stable over time and even task state (22), and individual differences in the location of whole-brain functional networks correlate with cognitive variability (23, 24). This raises a compelling hypothesis that the spatial organization of brain networks may reflect traits or characteristics of an individual, and may thus shed light on behavior in neurodevelopmental disorders such as autism. If the spatial organization of functional regions impacts behavior, we may expect to see the greatest disruptions in spatial organization of regions that are important for social or communicative functions most altered in autism.

### Why might functional neuroanatomy for faces be altered in autism?

Our finding of atypical spatial organization of the face network in autism complements a large literature on atypical structure and function of face-sensitive regions in the brain in autism. Functionally, regions like the fusiform face area are consistently under-active in ASD, show atypical response profiles, and are atypically functionally connected to other face processing regions when compared to NTs (e.g., (25, 26). Only a handful of studies have either directly or indirectly investigated the spatial organization of these regions, finding preliminary evidence that face processing does not occur in expected locations in autism (12, 13).

Autism provides a particularly compelling condition in which to examine the origins and consequences of variability in functional topography. In particular, autism has been associated with atypical patterns of early brain development, with some evidence suggesting disruptions emerging even in utero that may alter the initial layout of functional systems. These early perturbations could contribute to shifts in the spatial organization of cortical networks. Indeed, the emergence of functional regions is highly influenced by early genetic factors that initiate the process of patterning the cortex (27). Disruptions to genes important for patterning alter the size and organization of functional areas (28). A handful of genetic studies of ASD implicate several genes that guide the development and appropriate patterning of the cortex (see (29) for a review).

Importantly, autism is also characterized by substantial heterogeneity in sensory experience, which may further shape, or amplify, variation in the location and organization of functional regions over development. This is particularly important as both functional specialization and location of cortical regions relies on appropriate sensory input (30). In absence of typical sensory experiences, cortical regions may not emerge in optimal locations, with knock on effects on functional specialization and ultimately behavior. Unlike neurotypical individuals, for whom sensory experiences are relatively constrained by shared preferences, sensory preferences in autism are atypical and highly variable from a young age. For instance, while neurotypical infants tend to show a strong preference for the face and eyes, even moments after birth, infants with a high likelihood of developing autism often avoid eye-gaze and show diminished attention to social and face stimuli (10, 11). These behavioral differences emerge as early as 12 months (31). Variability in visual preferences in turn contributes to a more heterogeneous sensory experience for autistic children, which may in turn affect functional specialization of sensory areas. Here, we identified brain-wise displacement in how face processing is anchored to cortical topography in autism. While our participants are adults, such a shift likely reflects the outcome of the interplay between intrinsic developmental and extrinsic sensory processes that contributes to the functional topography of the mature cerebral cortex (e.g., (32)).

Other (non-autism) populations characterized by atypical sensory experiences support the idea that sensory experience influences the location and spatial variability of functional neuroanatomy. Increased variability in functional neuroanatomy has also been observed in sensory-*deprived* populations: congenitally blind and deaf individuals show greater variability in the connectivity patterns of primary and more widespread visual and auditory regions, respectively (33, 34). Blind individuals, who access reading via braille, show additional reading-specific functional regions in posterior parietal cortices important for touch, far from their traditional location in the visual word form area in occipitotemporal cortex (35). Similarly, the word-specific peak response is located farther from its canonical location in illiterates compared to individuals who learn to read in adulthood, and compared to typical readers who were exposed to print in early childhood (36, 37). This suggests that shared versus highly diverse sensory experiences may have opposite influences on the degree of spatial variability in the organization of functional regions. In many autistic individuals, for whom quality and quantity of face processing differs significantly from NT patterns, even quite early in development, it is perhaps unsurprising that face-sensitive regions are among the most noncanonically organized. At the same time, given the correlational nature of fMRI, these findings cannot adjudicate causality: it remains unclear whether altered spatial organization drives differences in face processing, reflects them, or emerges through reciprocal interactions over development.

### Regional differences in dispersion of individually defined functional regions in autism

Notably, the greatest dispersion in face-sensitive regions in autism was found in posterior occipito-parietal regions (including the fusiform face area). Increased heterogeneity in these occipito-parietal is particularly striking given that these regions usually show particularly *low* spatial variability in neurotypical brains (38). Posterior face-sensitive regions (such as the FFA and OFA) are typically activated earlier during face processing and primarily important for face identification, while frontal regions are engaged later for higher-order face-processing such as recognition or memory. Posterior occipito-parietal face regions also emerge and mature earlier in development (39), and are therefore less likely to be influenced by experiential factors. On the other hand, frontal face processing regions (e.g., mPFC) are typically associated with more abstract aspects of face processing (emotion, recognition), are located in parts of the brain that emerge later in development, and may thus be influenced by variability in inputs and experiences. The present finding of atypical functional neuroanatomy specific to posterior brain regions suggests that disruption to the organization of face processing regions likely occurs for earlier processing stages and perhaps earlier in development, and that this impacts social communication behaviors. This interpretation is in part supported by genetic studies of ASD finding greater gene expression differences in posterior regions (40).

Of note, dispersion, but not displacement, of functional regions was influenced by region (posterior vs. frontal). While there may be many explanations for this pattern, we suggest that the face network in autism is globally shifted from its canonical location in part due to increased intersubject heterogeneity in the precise spatial location of face-sensitive functional regions. Importantly, the topography of frontal lobe face regions was *generally* more heterogeneous across all individuals (both NTs and ASD individuals, see **Figure 2D**). The lack of group differences in dispersion in frontal regions may therefore reflect a “floor” effect across all participants, whereby all individuals show sufficient variability in these regions to make it difficult to identify any group differences. Atypical organization of occipito-parietal face regions specifically was associated with more atypical social communication in autism. No such relationship was detected for frontal regions. It could be the case that small differences in posterior occipito-parietal face regions (which form the basis of early face perception) may have large effects on behavior, especially given that these regions send inputs to prefrontal regions. It may also be the case that the patterning of posterior functional regions is more discretized than in the frontal lobe-with specific (yet proximal) regions engaged in the processing of distinct category types. Thus, small shifts in the precise location of posterior functional regions could result in greater disruptions to behavior.

### Atypical functional neuroanatomy in autism has broad implications for neuroimaging

Traditional neuroimaging analyses rely on averaging across individuals and reporting univariate group-level effects at each voxel. This approach assumes relatively strict voxel-wise correspondence in functional neuroanatomy across individuals and across groups, such that spatial averages respect central tendency (41). In populations such as autism, for whom functional neuroanatomy may be atypical and highly variable, group averages or the use of anatomically defined regions of interest may inaccurately represent or even underestimate brain responses. Under this framework, it is possible that findings of reduced face-related activation in autism could reflect reduced *overlap* of functional regions across individuals, rather than true differences in brain magnitude (illustrated in **Figure 5B**; see also (9)). Some of these issues can be addressed by precision fMRI approaches used in the present study, which identify the top activation for each individual within a relevant parcel and thus account for some degree of individual variation in brain organization. However, taking account of *both* group averaging and precision approaches could play an important role in uncovering mechanistic differences in functional neuroanatomy in autism. Indeed, in prior work examining a subset of the participants reported here, we observed reduced repetition suppression in the fusiform gyrus in autism relative to NTs - which may seem at odds with the findings described here. However, our prior work used *a priori* regions of interest, which were highly overlapping with the *expected* location of the fusiform face area in NTs, rather than individualized regions of interest (25). When defining face-sensitive regions *within individuals* in the present study, we found no group differences in repetition suppression magnitude. While the response magnitude in individually-defined face regions did not differ between groups, relying on functional regions that are not optimized for the work of face processing still had significant impacts on behavior (as seen here by significant correlations between autism symptoms and spatial displacement of the face network). Converging findings between the group- and precision-imaging approaches suggests that autistic individuals do not properly activate the expected or canonical location of the FFA (25), but rather rely on a more distributed set of regions for face processing. Both failure to activate traditional regions, and spatial displacement of individually-defined face regions have behavioral consequences. Together these results highlight the value of using multiple, complementary approaches to understand brain function in autism.

While precision fMRI allows for localization of individual functional regions, results must also be interpreted carefully, particularly within the context of clinical populations with high heterogeneity (like autism). For instance, even precision fMRI approaches could miss ‘true’ functional regions (and underestimate brain activation) if the parcels from which each individuals’ activation is extracted are not inclusive enough to capture the increased spatial variability observed in autism or other conditions. More broadly, assumptions about stability across runs are critical in precision fMRI approaches. In our data, we did not observe increased noise or reduced reliability across runs in ASD, suggesting that spatial variability is not simply a consequence of temporal instability. However, this may not generalize to other clinical populations, where greater noise or fluctuations in functional organization across runs could bias estimates. This could thus raise a methodological concern for precision approaches that define subject-specific functional regions in one run and extract responses in another.

### Limitations and future directions

These findings should be interpreted within the context of a few limitations. First, in the present study, we conducted secondary analyses of previously collected data, and the imaging task analyzed (repetition suppression) was not designed with consideration of the specific analyses used here. Repetition suppression is a classical task for identifying regions that are highly responsive and stimulus-specific (15), but additional studies defining face regions based on contrasts between stimulus categories (e.g., faces versus objects in the same run) may reveal additional insights (42). To better understand this aspect of brain organization, future work should employ methods to investigate how the organization of selective voxels may vary for autistic individuals. This line of inquiry can better inform the potential mechanisms underlying the atypical organization found in the present study.

Additionally, because participants were adults, it is not possible to determine the extent to which differences in functional organization reflect variation in developmental history, including the type and timing of interventions. Given the long developmental interval, experiential factors may have shaped the organization of these systems over time, limiting our ability to infer their origins. Future work should probe the developmental timeline and mechanisms underlying the more variable organization of functional regions in the brain in autism. Given the lifelong pattern of atypical sensory experiences in autism spectrum disorder, it is possible that the organization of functional regions becomes *more* variable over time. A longitudinal study that examines the organization of functional regions throughout childhood would help to clarify when this variability emerges and changes over time. Additionally, it is important to further investigate the functional consequences of a more heterogeneous organization of functional neuroanatomy. What are the functional consequences if an individual has the same magnitude of activation but with a less typical spatial organization? Of course, there is also a possibility that this atypical organization could result in processing strengths in certain domains. Indeed, there is some evidence that FFA activation is similar to, or greater than, that of neurotypical individuals when probed with stimuli in which autistic individuals have particular expertise or interest (e.g., special interests, mother’s face; (43–45); But see (5) about whether FFA is related to visual expertise or face processing. Finally, there is increasing evidence for dimensions of variability within autism, such as individuals receiving diagnoses at younger or older ages (46). A careful consideration of this variability in autism could refine our interpretations of prior studies that reported reduced activation in certain brain areas and help to reconcile inconsistent findings in the field.

A final concern is our focus on autism. Future work should explore whether the increased variability identified in here is specific to autism spectrum disorder, or more general to neurodevelopmental conditions broadly. Moreover, many neuroimaging studies involve groups comparisons, such as between clinical and control groups or between developmental age groups. Nearly all of these studies that report differences use group averaging of activations in a common selected space. The present findings suggest that some of these studies may reflect not less activation in a group, but rather more diverse loci of activations in one group relative to another group. Future imaging studies, in clinical and developmental populations, ought to consider both possibilities in order to interpret group differences accurately.

## Materials and Methods

### Participants

Adults (age 18-45 years) were recruited from the greater Boston area of the United States on the basis of their self-reporting a prior diagnosis of autism (ASD group) or not (neurotypical, NT, group). All participants completed the *Kaufman Brief Intelligence Test* (*KBIT*) Matrices subtest as a measure of general cognitive ability (47). Participants also completed the *Autism Quotient* (*AQ*; (48)) as a measure of autistic traits (Table 1). Autistic participants’ self-reported autism diagnosis was confirmed by research-reliable administration of the Autism Diagnostic Observation Schedule - 2 (ADOS-2; (49)). Exclusion criteria across both groups included history of hearing or visual impairments and incidental findings on MRI. NT participants were excluded if they reported a family history of autism or other neurological condition.

**Table 1.**
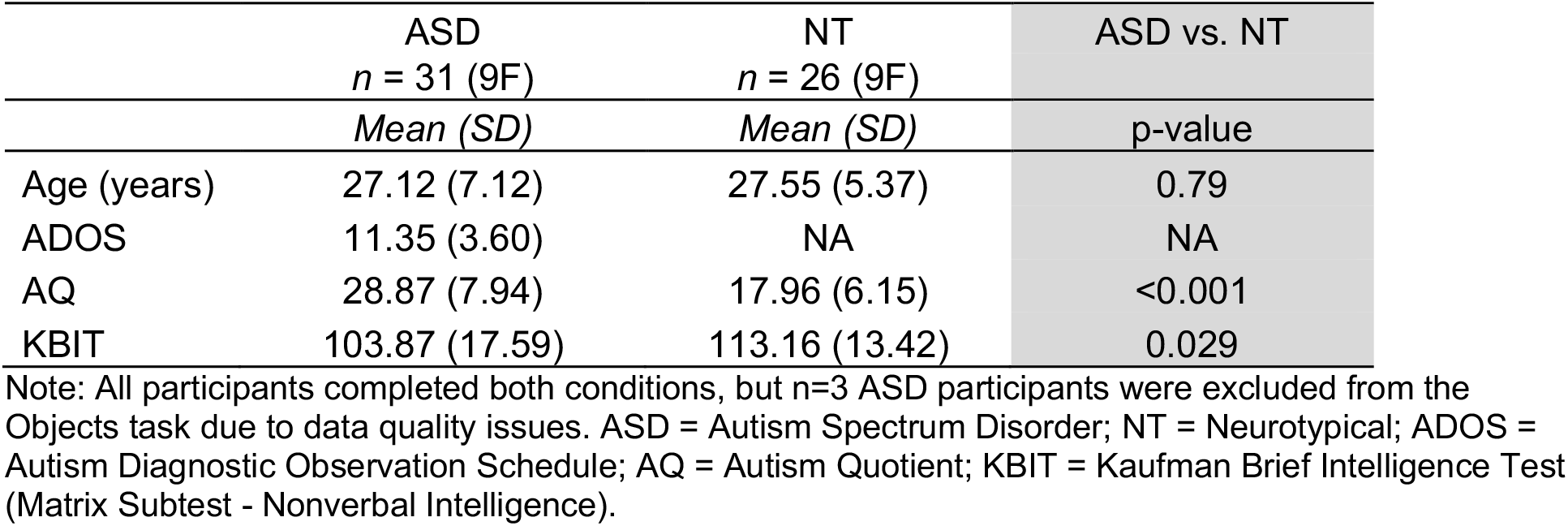
Participant demographic information and behavioral assessment scores by group and task.

|  | ASD<br>*n* = 31 (9F) | NT<br>*n* = 26 (9F) | ASD vs. NT |
| --- | --- | --- | --- |
|  | *Mean (SD)* | *Mean (SD)* | p-value |
| Age (years) | 27.12 (7.12) | 27.55 (5.37) | 0.79 |
| ADOS | 11.35 (3.60) | NA | NA |
| AQ | 28.87 (7.94) | 17.96 (6.15) | <0.001 |
| KBIT | 103.87 (17.59) | 113.16 (13.42) | 0.029 |

Note: All participants completed both conditions, but *n*=3 ASD participants were excluded from the Objects task due to data quality issues. ASD = Autism Spectrum Disorder; NT = Neurotypical; ADOS = Autism Diagnostic Observation Schedule; AQ = Autism Quotient; KBIT = Kaufman Brief Intelligence Test (Matrix Subtest - Nonverbal Intelligence).

Among 68 recruited participants (ASD: *n* = 39, NT: *n* = 29; as reported in (25)), *N* = 58 were eligible and provided sufficient data for inclusion in the present study (ASD group: *n* = 31; 9 female [sex], 21 male; *M*_age_= 27.5 years, *SD*_age_ = 7.3; NT group: *n* = 26; 9 female, 17 male; *M*_age_= 27.6, *SD*_age_ = 5.4).

All participants provided written informed consent and were compensated for their time in accordance with the Massachusetts Institute of Technology (MIT) Committee on Use of Humans as Experimental Subjects (MIT’s Institutional Review Board).

### Experimental design

#### FMRI task

The goal of the present study was to characterize differences in the spatial organization of stimulus-specific functional regions across autistic and non-autistic adults. To do this, we used a series of classical ‘repetition suppression’ fMRI tasks that robustly identify and engage stimulus-specific functional regions (50). Data for the present study come from a larger study that investigated repetition suppression across multiple stimulus categories (including faces probed here; (25)). Participants passively viewed blocks of repeating and non-repeating faces, interspersed with rest blocks (9.6s per block; total of 10 blocks of each per run of each task), while in the scanner. Task order was counterbalanced across participants. In “repeat” blocks, a single face was repeated eight times. In “non-repeat” blocks, eight unique faces were presented. Faces were not repeated across blocks, and remained on the screen for 700ms, with a 500ms inter-trial interval. Rest blocks were the same length as task blocks, and participants fixated on a “+” and waited for the next face to appear. Within each stimulus category, a total of 160 unique stimuli were assigned to the *non-repeat* condition, and 20 unique stimuli were assigned to the *repeat* condition. In each block, as a measure of task engagement, participants were required to press a button in response to infrequent target stimuli, which were denoted by vertically inverting a single random image in each block. Stimuli were presented in the center of an 1024×768 pixel display with a dark gray background. Face stimuli consisted of 180 grayscale photographs of male and female individuals with neutral expressions. Each photograph was cropped close to the face to minimize hair and background (size: 256×256 pixels). For additional paradigm details see (25) and (16). Hit rate for infrequent targets (number of hits/number of targets) did not differ between groups (NT = 96.7%, ASD = 95.4%, *p* = .60).

#### FMRI acquisition

Data were acquired on a Siemens PRISMA 3T scanner with a 32-channel head coil. At the beginning of each scanning session, a high-resolution T1-weighted (T1w) multi-echo MPRAGE volume was acquired (TR = 2530ms, TE = {1.69, 3.55, 5.41, 7.27ms}, voxel size = 1.0mm^3^, slices = 176, FOV = 256mm^3^). T2*-weighted EPI functional scans were collected using simultaneous multi-slice acquisition (interleaved, acceleration factor = 5) and contained 348 volumes per run (total of 696 volumes per task; TR = 850ms, TE = 32ms, voxel size = 2.5mm^3^, slices = 55, FOV = 210mm^3^).

### Statistical analysis

#### FMRI preprocessing

Preprocessing was performed using fMRIPrep. Each T1w volume was corrected for intensity non-uniformity and skull-stripped. Spatial normalization to the ICBM 152 Nonlinear Asymmetrical template version 2009c was performed through nonlinear registration with ANTs v2.1.0, using brain-extracted versions of both the T1w volume and template. Brain tissue segmentation of cerebrospinal fluid (CSF), white-matter (WM) and gray-matter (GM) was performed on the brain-extracted T1w using FSL *fast*. Functional data were motion corrected using FSL *mcflirt* and co-registered to the T1w image using boundary-based registration with FSL *flirt*. The motion correcting transformations, BOLD-to-T1w transformation, and T1w-to-template (MNI) warp were concatenated and applied in a single step using ANTs. Normalized functional images were smoothed with a 6mm FWHM Gaussian kernel to reduce uncorrelated spatial noise. Framewise displacement was calculated for each functional run. Outliers were defined as any volume for which framewise displacement was > 0.5mm. Participants with > 20% of volumes marked as outliers were excluded from further analysis for that task.

#### FMRI modeling and analysis

All first- and second-level modeling was conducted in SPM12. Normalized and smoothed images were entered into a first-level general linear model. For each run, repeating and non-repeating blocks were entered as regressors. Outliers were entered as nuisance regressors. Contrasts for repetition suppression (non-repeat > repeat) were estimated. These activation maps, representing regions that are sensitive to a particular stimulus category (50) were then used to probe differences in the spatial organization of functional neuroanatomy between individuals and across groups.

#### Creation of group-constrained parcels

Because the precise location of functional regions in a standard space varies across individuals, we employed data-driven ‘precision fMRI’ methods that identify individual defined regions of interest in each participant’s brain, rather than imputing regions based on population-level assumptions regarding macroanatomical landmarks, published coordinates, or group averages. In this group-constrained, subject-specific approach (GCSS; (18)), we first identify large regions (‘parcels’) that capture the probabilistic extent of activation across participants. These parcels were used to then identify each participant’s individually defined regions (see *Identification of Participant-Specific Functional Neuroanatomy* below). Parcels were defined using data from NT participants, consistent with prior literature applying precision methods to clinical groups (e.g., (51)). To create the parcels, each NT participant’s statistical parametric map of the repetition suppression effect (non-repeating > repeating) was thresholded voxel-wise at *p*<.001 (uncorrected) and binarized. The binarized maps from these participants were overlaid to create a probability map for the NT group which was smoothed at 6mm FWHM and thresholded voxel-wise at *n*=2 subjects. A watershed algorithm from the SPM-SS toolbox was used to detect and create volumes around local probability maxima, thereby parcellating the probability map. Parcels which contained significant voxels from 80% or more of NT participants were retained for analysis. Thus, because the parcels represent relatively broad but contiguous regions in which a large percentage of participants show significant activation, they can be used to constrain the selection of participant-specific regions which, though not necessarily in the same exact location in standard space, are nonetheless functionally homologous across individuals. For faces, we identified 5 parcels that correspond to known regions of the face processing network, including parcels encompassing the medial prefrontal cortex, the right middle frontal cortex, the right posterior parietal lobe, and bilateral occipito-temporal cortex (including the fusiform gyrus). All subsequent participant-level analyses were conducted within these parcels.

#### Identification of Participant-Specific Functional Neuroanatomy

For each participant (in the NT and ASD groups), we identified the top 10% of activated voxels within each face-processing parcel (i.e., each participant’s functional region of interest). This parcel-constrained voxel selection approach allows for substantial inter-individual variability in the precise spatial location of brain responses. This approach ensured that we captured each participant’s own most face-specific functional regions and that these regions were the same size across participants. Although most analyses used the top 10% of activated voxels to identify functional regions, we also defined functional regions at additional thresholds (top 20%, 30%, 40%) to look more liberally at face-specific functional neuroanatomy.

#### Spatial organization of functional regions

We conducted two separate analyses to characterize the spatial organization of functional regions within and across diagnostic groups: *displacement* and *dispersion*. Notably, stimulus sensitive regions are distributed throughout the brain, and different regions are thought to have slightly different functions. Given this, and evidence for a temporal hierarchy of processing from posterior to frontal regions (which typically engage in higher-level stimulus processing), we also assessed the effects of global location (posterior, frontal) on group differences in functional neuroanatomy.

#### Displacement of functional regions

We examined the distance between each participants’ face region and a functionally defined group-level centroid of activation based on the NT group. We used this measure to quantify how ‘far’ participants’ face regions were located relative to their expected location in the brain (typically highly overlapping in NTs). To determine this anatomical distance, we first extracted the MNI-space anatomical coordinates (i.e., X, Y, Z) of each voxel in each participants’ functional region of interest, by parcel. To identify the functional center of each parcel, we calculated the centroid of the NT participants’ coordinates of activation (See **SI-Table 1** for coordinates). Then, in each parcel, we calculated the Euclidean distance between each voxel in a participant’s functional region of interest and the centroid for that parcel. We categorized the parcel location as either posterior (i.e., fusiform face area, occipital face area, and posterior parietal cortex) or frontal (i.e., mPFC and inferior frontal gyrus).

A linear mixed effects model was used to assess effects of *group* (ASD, NT) and location (anterior, posterior) on Displacement. The following model was used:

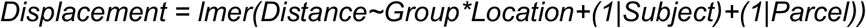

#### Dispersion of functional regions

We also quantified the degree of spatial overlap between each participant’s face regions and those of all the other participants in their group. This measure allowed us to investigate how similar the location of functional regions are at the voxel-level across participants and then compute group differences in average overlap. To do this, we used the Jaccard Index (52), a measure of the extent to which two activation maps overlap (53–55). We calculated the Jaccard Index for each participant pair within each group as the proportion of voxels in common between the two maps out of the total voxels across both of the maps:

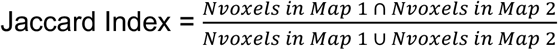

Linear mixed effects models were used to assess effects of *group* (ASD, NT) and location (anterior, posterior) on Jaccard Index. The following model was used:

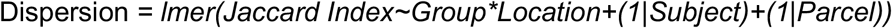

Additionally, we examined whether the size of our individually defined regions was influencing our results within the posterior region. For instance, it is possible that defining face regions as the top 10% of activated voxels did not sufficiently capture the extent of each autistic individual’s face-sensitive areas, or that functional neuroanatomy is generally less “peaky” and more distributed in autism than in NTs. We compared the Jaccard Index for three increasingly liberal thresholds (top 20%, 30%, 40% of most activated voxels), resulting in progressively larger individually defined regions. We restricted these analyses to the posterior face regions, based on the identified group differences in these regions.

#### Correlation between displacement of functional regions and behavior

We investigated how displacement in functional neuroanatomy may parallel differences in behavior. To do this, we conducted a one-tailed Spearman’s Rank correlation between each ASD participant’s mean Euclidean distance (i.e., displacement from the group; described above) and ADOS total score (a standardized diagnostic measure of autistic behavior and symptom severity). Notably, we did not examine the correlation between dispersion and behavior, as the dispersion reflects inter-individual variability in the spatial organization of functional regions and is therefore a group-level property, making it less meaningful for direct correlation with individual behavioral measures. In contrast, the displacement measure provides a well-defined, continuous measure at the individual level, quantifying how far each participant’s functional regions deviate from a canonical location.

#### Investigating category-selectivity in group differences in functional neuroanatomy

To investigate whether identified differences in functional neuroanatomy were specific to face processing, we repeated the analyses of “Displacement” and “Dispersion” (as described above) on data from a repetition suppression task with object stimuli. The task parameters were identical to the face repetition suppression task, and participants completed 2 runs of the object task in the same fMRI session as faces (Described previously in (25)). Object stimuli consisted of 180 color photographs of objects in isolation on a white background (size: 256×256 pixels).

Using the same GCSS approach as described above, we created 6 object-sensitive parcels including bilateral middle frontal cortex, left posterior parietal cortex, bilateral occipito-temporal cortex, and lateral occipital cortex (**SI-Figure 1**). These object-sensitive parcels were used to extract object-specific fROIs in each participant and assess whether group differences were category specific.

#### Control analyses of magnitude and selectivity

Finally, we conducted a series of control analyses to examine alternative explanations for the results. We first examined the noisiness of fMRI data (Spearman’s Rank correlation of voxel-wise fMRI magnitude between runs within individually defined functional regions from each participant).

We next assessed whether there were group differences in overall magnitude of the adaptation contrast within individually defined functional regions. For instance, if autistic individuals did not activate certain regions at all, this would by definition reduce measures of overlap in functional regions across participants. For the magnitude analyses, the following model was used: lmer(magnitude∼parcel*Group + (1|subID)).

Finally, we examined whether there were group differences in selectivity of responses to objects in the face fROIs or to faces in the object fROIs. Using each participant’s fROIs from Run 1 (Faces), we extracted the magnitude of non-adapt activation from Run 2 from both tasks (Faces and Objects). For each participant, we found the difference in activation between Faces and Objects for each parcel (mean difference score) as a measure of selectivity and compared these between groups.

## Supplementary Information for

Neurodiversity in the brain: More variable localization of face regions in autism

**SI-Figure 1.**
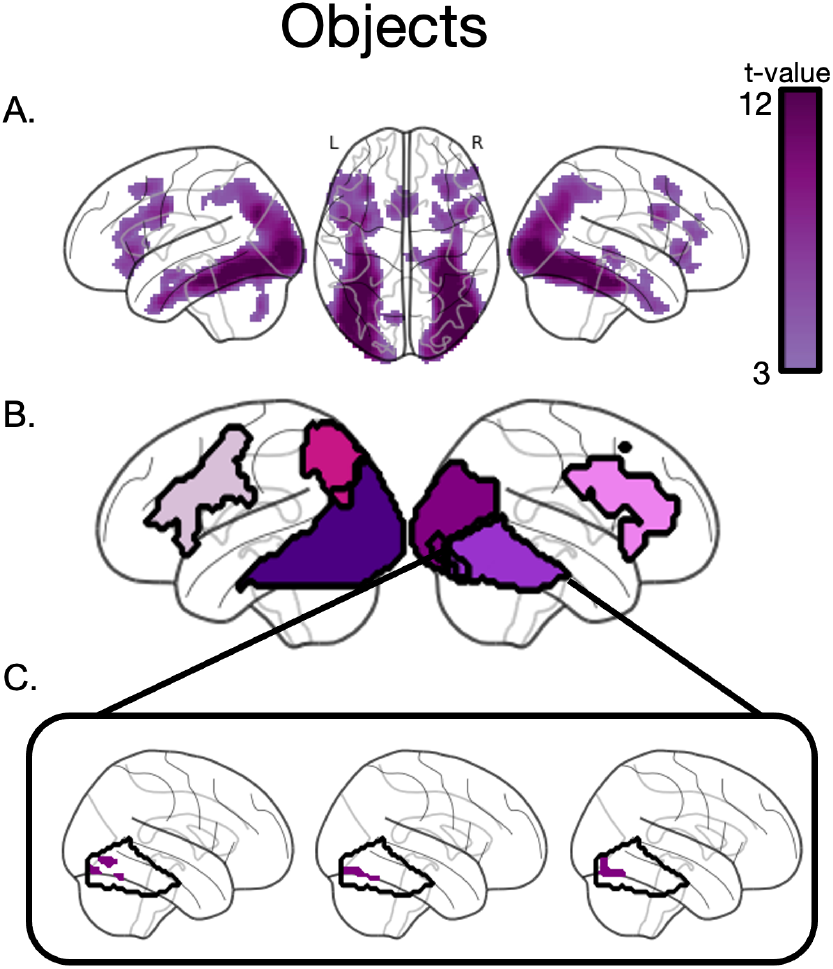
Object functional tasks activate object-specific functional regions. **A**. Group-level activation across all participants for objects, thresholded at *p*<0.001, FWE-corrected. **B**. Six functional data-driven parcels were identified that capture areas of high probability of object-specific activation across participants (Methods 3.3). **C**. Individually-defined functional face regions (top 10% of voxels within the parcel) from three example participants for the parcel including the FFA (Methods 3.4).

**SI-Table 1.**
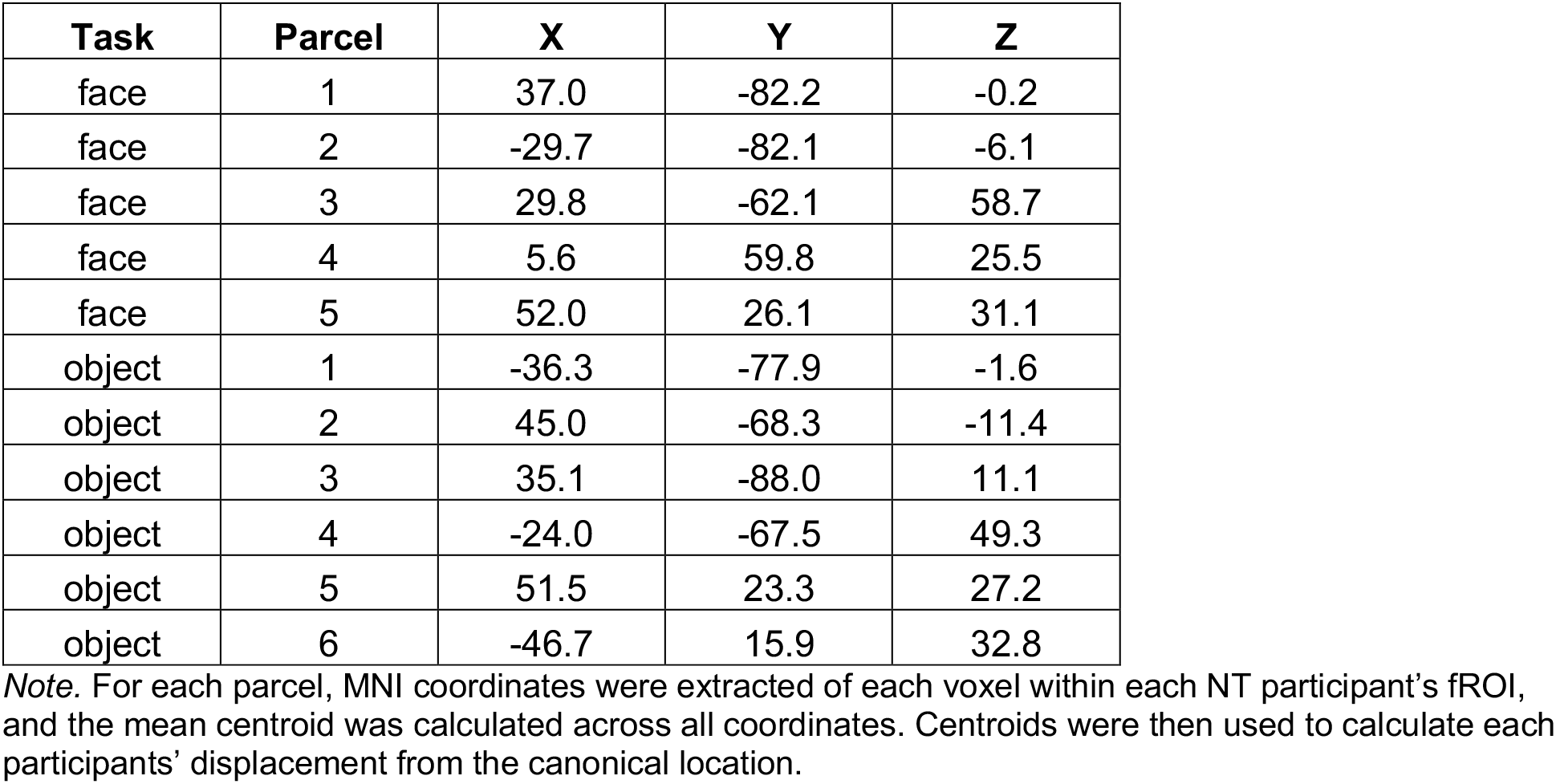
Canonical locations for each parcel, based on centroid of neurotypical activation.

**SI-Table 2.**
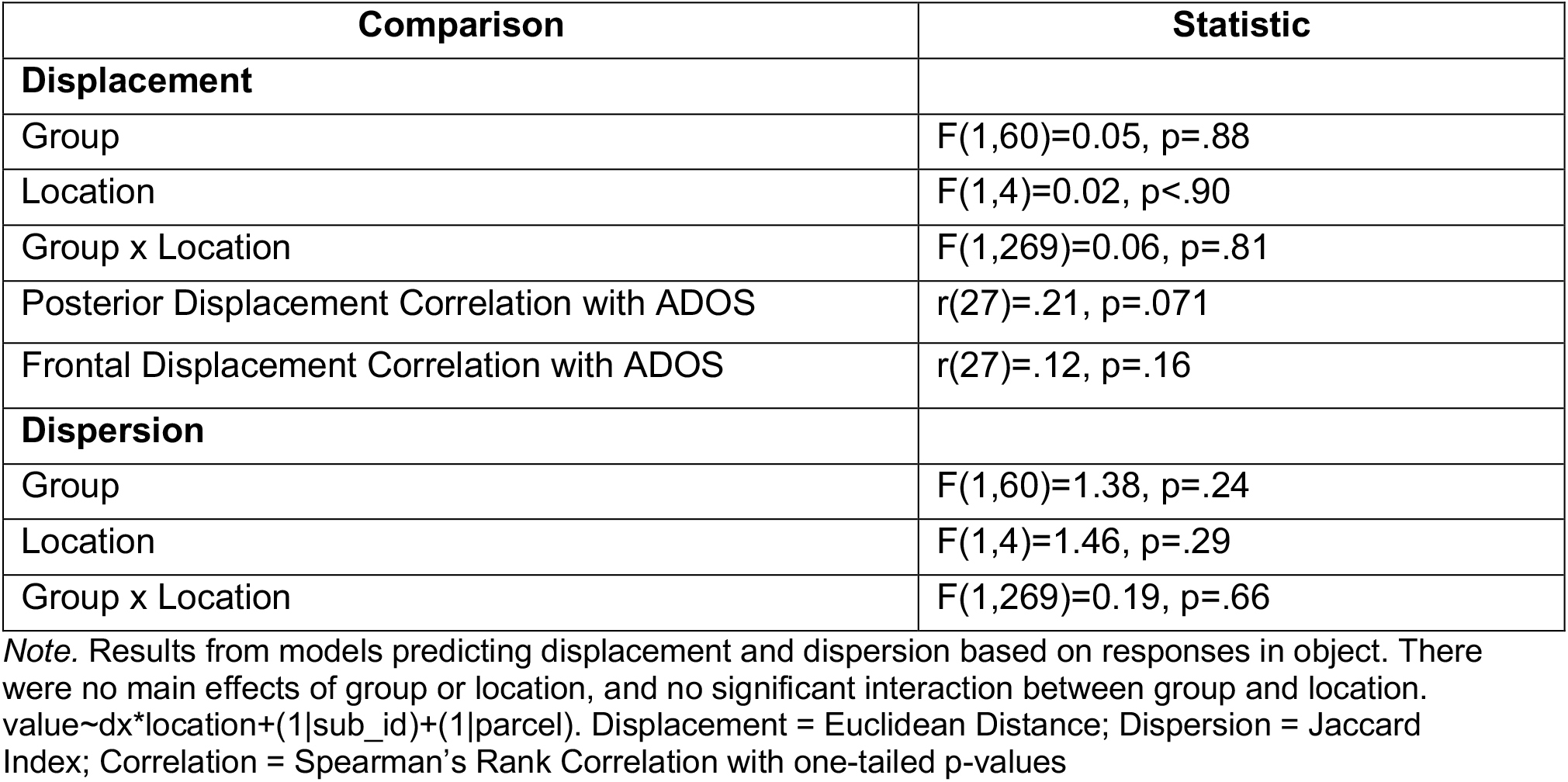
Displacement and Dispersion Analyses of Object-Specific Activation.

