## Supplementary material for "Neurodiversity in the brain: More variable localization of face regions in autism": SI materials

**Please note:** This is a preprint. It has not yet been peer reviewed.

**Supplementary Information for:**
Neurodiversity in the brain: More variable localization of face regions in autism

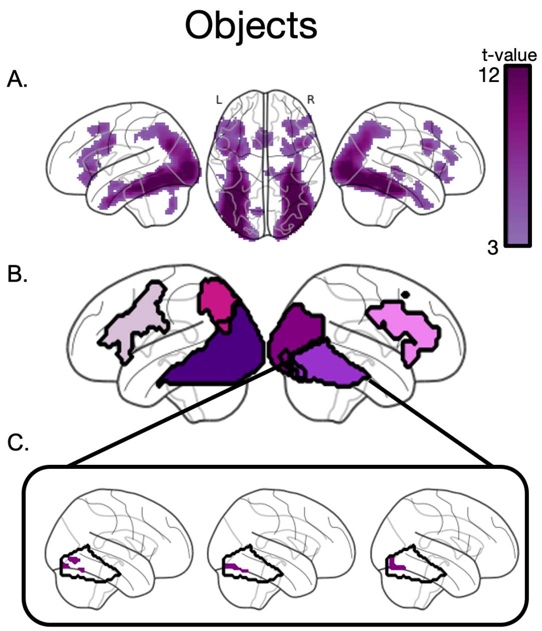


**SI-Figure 1.** Object functional tasks activate object-specific functional regions. **A.** Group-level activation across all participants for objects, thresholded at *p*<0.001, FWE-corrected. **B.** Six functional data-driven parcels were identified that capture areas of high probability of object-specific activation across participants (Methods 3.3). **C.** Individually-defined functional face regions (top 10% of voxels within the parcel) from three example participants for the parcel including the FFA (Methods 3.4).

**SI-Table 1**

Canonical locations for each parcel, based on centroid of neurotypical activation

| Task | Parcel | X | Y | Z |
|---|---|---|---|---|
| face | 1 | 37.0 | -82.2 | -0.2 |
| face | 2 | -29.7 | -82.1 | -6.1 |
| face | 3 | 29.8 | -62.1 | 58.7 |
| face | 4 | 5.6 | 59.8 | 25.5 |
| face | 5 | 52.0 | 26.1 | 31.1 |
| object | 1 | -36.3 | -77.9 | -1.6 |
| object | 2 | 45.0 | -68.3 | -11.4 |
| object | 3 | 35.1 | -88.0 | 11.1 |
| object | 4 | -24.0 | -67.5 | 49.3 |
| object | 5 | 51.5 | 23.3 | 27.2 |
| object | 6 | -46.7 | 15.9 | 32.8 |

*Note.* For each parcel, MNI coordinates were extracted of each voxel within each NT participant's fROI, and the mean centroid was calculated across all coordinates. Centroids were then used to calculate each participants' displacement from the canonical location.

**Please note:** This is a preprint. It has not yet been peer reviewed.

**SI-Table 2.**

Displacement and Dispersion Analyses of Object-Specific Activation

| Comparison | Statistic |
|---|---|
| **Displacement** | |
| Group | F(1,60)=0.05, p=.88 |
| Location | F(1,4)=0.02, p<.90 |
| Group x Location | F(1,269)=0.06, p=.81 |
| Posterior Displacement Correlation with ADOS | r(27)=.21, p=.071 |
| Frontal Displacement Correlation with ADOS | r(27)=.12, p=.16 |
| **Dispersion** | |
| Group | F(1,60)=1.38, p=.24 |
| Location | F(1,4)=1.46, p=.29 |
| Group x Location | F(1,269)=0.19, p=.66 |

*Note.* Results from models predicting displacement and dispersion based on responses in object. There were no main effects of group or location, and no significant interaction between group and location. value~dx*location+(1|sub_id)+(1|parcel). Displacement = Euclidean Distance; Dispersion = Jaccard Index; Correlation = Spearman's Rank Correlation with one-tailed p-values
